# Thrifty phenotypes in ants: Extending a human developmental hypothesis to a superorganism

**DOI:** 10.1101/2024.09.29.615703

**Authors:** Érik Plante, Ehab Abouheif, Jean-Philippe Lessard

## Abstract

Adaptation to extreme environments is commonly assumed to occur through microevolutionary change. However, studies of high-altitude human populations show that organisms can also respond to nutritional stress through developmental plasticity, producing thrifty phenotypes that prioritize essential function over costly structures. Whether this adaptive strategy extends beyond humans, and how it operates in natural systems, remains largely unexplored. Here, we test the thrifty phenotype hypothesis (TPH) in a widely distributed superorganism, the carpenter ant *Camponotus herculeanus*, which inhabits some of the most environmentally challenging regions of the Northern Hemisphere. Colonies comprise inexpensive minor workers and energetically costly major workers, providing a powerful system for examining plastic investment under resource limitation. We quantified relationships between caste structure and climate and conducted a common-garden experiment to test the TPH. We show that the proportion of major workers declines with increasing latitude, independently of body size and colony size, and is best predicted by the number of days, annually, during which workers can nurse brood. Experimental results further demonstrate that colonies rapidly and plastically adjust caste structure in response to environmental conditions. Our findings reveal that thrifty phenotypes can emerge in superorganisms and could represent a conserved developmental response to environmental stress. By extending a central hypothesis from human biology to social insects, this work provides a unifying framework for understanding how developmental plasticity shapes adaptation across levels of biological organization.

## Introduction

Adaptation can be challenging in stressful environments where both energy and key nutrients are chronically limited. The evolutionary and developmental processes enabling organisms to adapt to stressful environments remain poorly understood [1–4]. Organisms may respond through evolutionary changes [5,6], phenotypic plasticity [7–9], or a combination of both. Populations of humans and other organisms inhabiting high altitude mountains or polar latitudes have been the focus of research on adaptation to extremely stressful environments[10–14]. Studies on high-altitude human populations, such as those of the Andean and Himalayan mountains [15,16], have revealed that nutritional stress during development can induce plastic adjustments consistent with thrifty phenotypes [17–20]. The *thrifty phenotype hypothesis* (TPH) [17] posits that nutrient shortages during development lead to selective investment in organs, where the growth of critical organs (e.g. heart, brain) is preserved at the expense of peripheral growth, resulting in shortened limbs (e.g., reduced ulna and limb length) relative to overall body size [15,16,21]. However, it remains unclear whether non-human organisms evolve thrifty phenotypes to adapt to stressful environments, and which external factors may drive their expression [22,23].

The TPH was originally proposed in medical literature to explain type 2 diabetes through the lens of developmental plasticity [17]. Malnutrition during fetal development can alter growth patterns and metabolism [17,24–26,9,27,28,18,29–31]. When such individuals experience abundant nutrition later in life, the thrifty phenotype may become maladaptive, increasing the risk of type 2 diabetes and other health problems, including obesity and cardiovascular disease [17,24,25,9,27–31]. However, when environmental conditions are persistently stressful throughout an individual’s entire lifetime, such as at high altitudes or latitudes, then thrifty phenotypes are predicted to provide an adaptive advantage. In such stressful environments, individuals must store sufficient energy during the growing season to survive long winters [22]. However, identifying environmental drivers of thrifty phenotypes in humans is complicated by long developmental periods and cultural buffering that obscure environmental effects [32].

The TPH highlights the existence of a sensitive period or window during development, during which environmental conditions are sensed, and organisms plastically adjust organ and tissue growth to match the perceived environment [22]. Adaptive developmental plasticity is a core principle of the field of ecological evolutionary developmental biology (eco-evo-devo) [33,34]. Although the TPH has been tested only in humans, eco-evo-devo predicts that such developmentally sensitive periods may be more conserved than might be expected, and may therefore apply more broadly across animals [35]. This prediction is based on the existence of a highly conserved developmental toolkit that is shared across metazoan animals, in which a single genome produces differentiated cell types, tissues, and organs in response to internal and external cues [36,37]. Extending the TPH beyond humans therefore predicts the evolution of a sensitive developmental window across animals that detects nutrient limitation and reallocates resources toward essential organs at the expense of others [38,39]. Thus, in eco-evo-devo terms, the TPH can be reframed as the expression of conserved developmental windows through which external cues canalize growth trajectories, generating thrifty phenotypes across a wider range of organisms [40].

Eusocial colonies provide a natural system to test the TPH, because a single genome produces morphologically distinct individuals or castes in response to environmental cues [41–43], making colonies superorganisms with traits analogous to organ-level traits in multicellular organisms [44,45]. Moreover, colony-level phenotypes, including caste structure, are directly observable [38,39]. Colony size (number of individuals) and colony structure (caste, age, or size distribution) are two phenotypic traits with adaptive value [46–48] that vary geographically [49–51], and respond to resource availability [41]. Although most species have a monomorphic worker caste, meaning that workers show little variation in size or head-to-body scaling, several species have evolved worker polymorphism, in which workers show dramatic variation in size and head-to-body scaling [51]. Worker polymorphism is therefore particularly informative because it reflects developmental allocation decisions in response to environmental conditions [50,51].

Broadly distributed species may encounter extremely stressful conditions at the edge of their range, imposing selective pressures on those edge populations that are distinct from those of core populations [52–55]. We therefore test the *thrifty phenotype hypothesis* (TPH) by investigating phenotypic variation among populations of a widely distributed superorganism, the carpenter ant *Camponotus herculeanus* (Linnaeus, 1758), whose northern range limit occurs in extremely stressful environmental conditions across the northern hemisphere. Eusocial colonies of *C. herculeanus* have a worker caste consisting of thousands of small-headed minor workers specializing in nursing and foraging, and larger-headed, more energetically costly, major workers specializing in defense and food storage [56]. Subcaste identity is determined by a developmental switch mediated by Juvenile Hormone (JH) during the last larval instar [57,58]. Variation in the proportion of majors and minors results from this developmental reprogramming, reflecting larval allocation decisions and can be influenced by environmental conditions. The TPH, when applied to eusocial organisms, predicts that colonies experiencing resource limitation should maintain overall colony size by plastically reducing investment in costly individuals, such as majors.

The TPH has been tested on human populations and via lab experiments on rodents [59,60]. Here, we performed the first test of TPH in natural systems outside humans by using whole-colony sampling across the temperate and boreal forest biomes of eastern Canada—by felling nesting trees and collecting all individuals—and relating colony worker caste structure to variation in micro- and macro-climate. Specifically, we collected 26 whole colonies of *C. herculeanus* distributed along an 8° latitudinal gradient from temperate forest to the northern edge of the boreal forest in eastern Canada (figure 2). This range encompasses both the core and northern edge of the species’ distribution. We quantified colony size, worker body size and caste structure, and estimated local environmental variables at each colony. To determine whether caste structure is evolutionarily fixed or plastic, five colonies from the core and four colonies from the northern edge were transplanted into a common rearing environment and assessed after one larval generation. Together, these approaches provide the first comprehensive test of the TPH in a non-vertebrate, offering insights into how developmental plasticity shapes colony-level adaptation under extreme environments.

**Figure 1.**
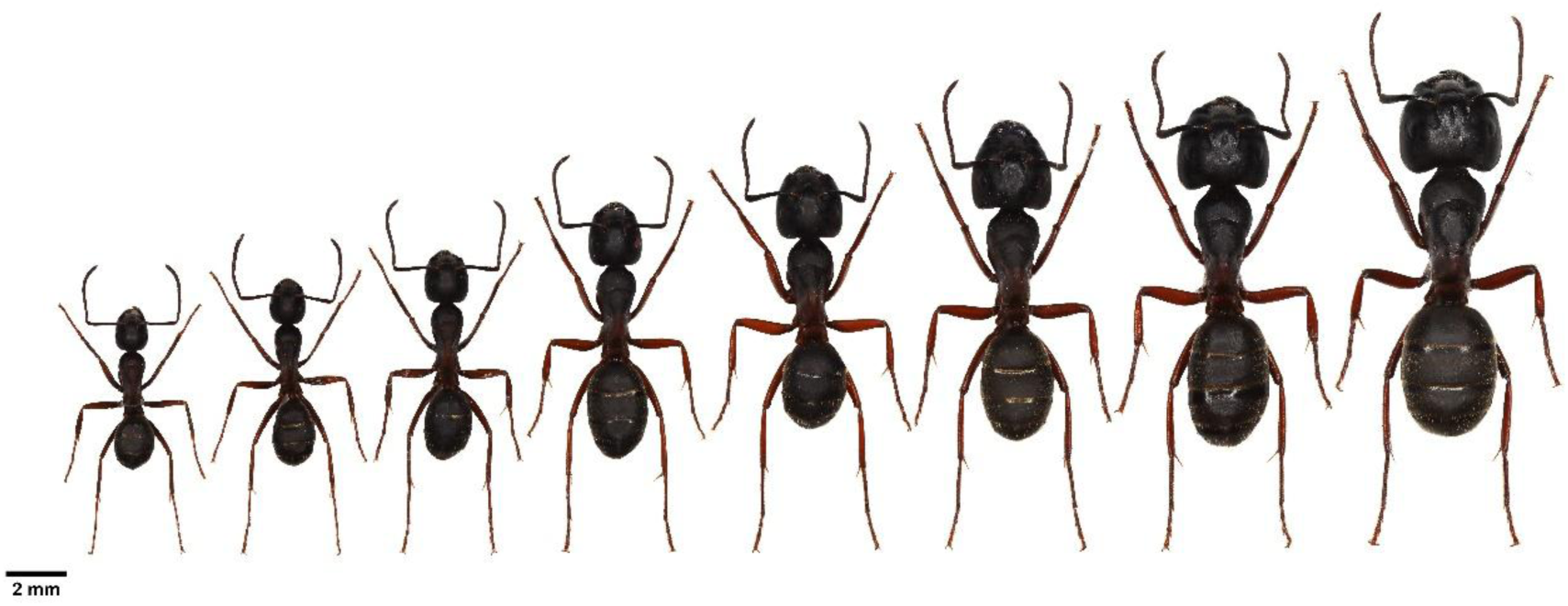
*Camponotus herculeanus* worker polymorphism (left-to-right) from minors to medias and majors. All workers originate from the same colony and were photographed at the same scale. These specimens represent the possible worker morphological range for a given large colony of the species in the study sites in the province of Quebec, Canada.

**Figure 2.**
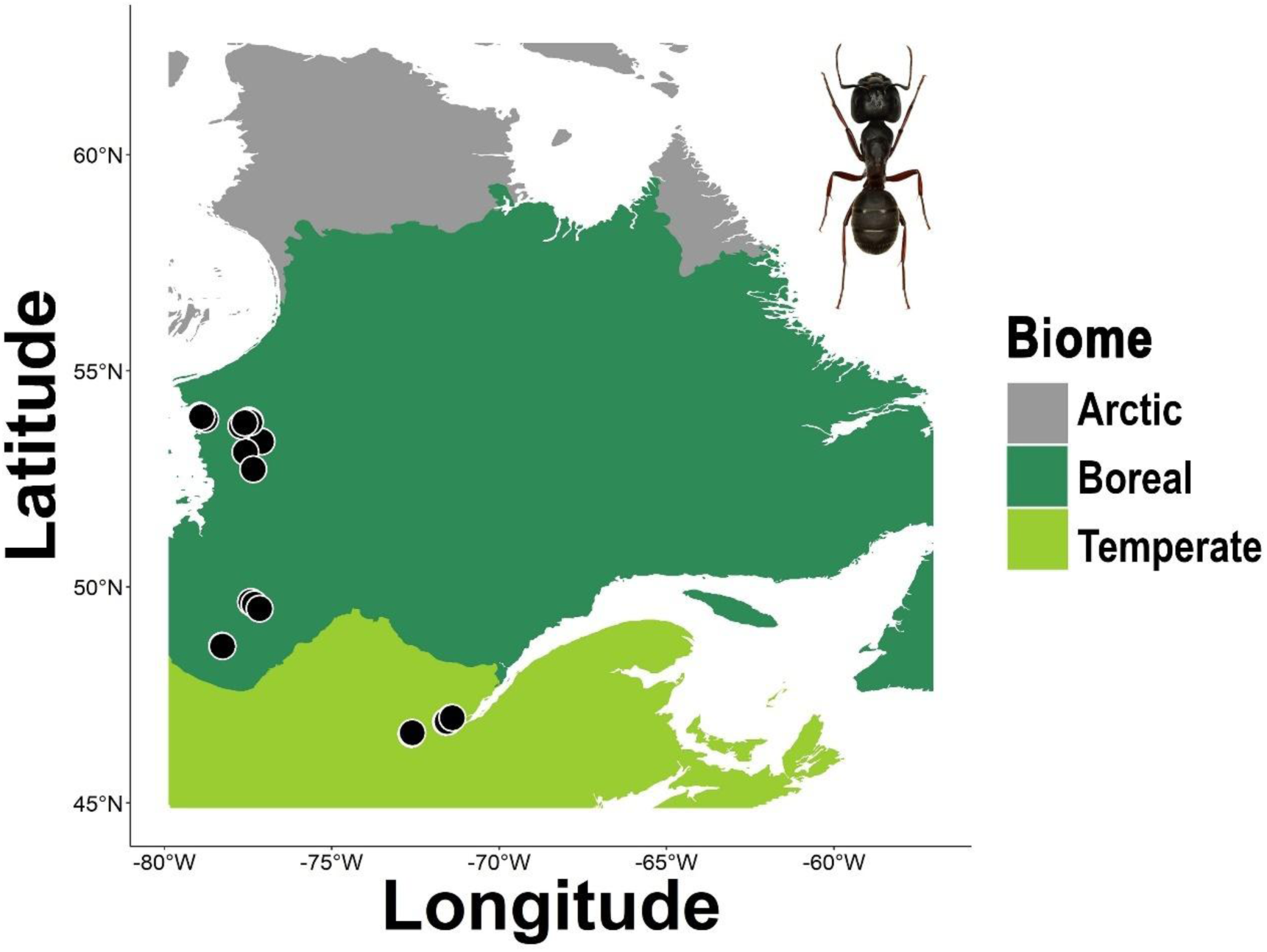
Location (*n* = 26) of the *Camponotus herculeanus* colonies used for the observational study in Eastern Canada (province of Quebec) across biomes [139].

## Results

The TPH predicts a reduced investment in costly major workers (i.e., reduced subcaste ratio) in stressful environments. Our data support the TPH, such that colonies in colder, drier environments invested less in major workers (electronic supplementary material, figure S1, table S4), whereas mean major worker body (electronic supplementary material, figure S1, table S4) and colony size (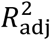= -0.04, df = 24, *F* < 0.01, *p* = 0.983) were not related to regional climate, indicating that colonies maintain consistent body and colony size regardless of environmental stress.

To test whether the thrifty colony phenotype arises from developmental plasticity or from microevolutionary adaptation, we reared colonies from the southern and northern ends of the gradient for one generation in a common controlled environment and compared caste structure before and after rearing. Microevolutionary adaptation would predict that colony traits are genetically fixed and remain unchanged under uniform conditions.

After one generation, southern colonies showed no significant change in caste structure (*t* = –0.31, df = 4.89, *p* = 0.77; figure 5*a*), whereas northern colonies showed a significant increase in the proportion of major workers (*t* = –2.82, df = 4.14, *p* = 0.045; figure 5*a*). These results indicate that the thrifty colony phenotype in northern environments is driven by developmental plasticity. Moreover, after rearing, northern and southern colonies converged in caste structure (*t* = 0.95, df = 4.06, *p* = 0.40), indicating plastic adjustment under uniform environmental conditions. In contrast, mean major worker body size did not change in either southern (*t* = –0.99, df = 7.57, *p* = 0.35; figure 5*b*) or northern colonies (*t* = 0.06, df = 4.10, *p* = 0.95; figure *5b*), consistent with the earlier results that body size is not environmentally plastic.

Having established that the thrifty phenotype arises from plasticity, we next examined which environmental factors best predict caste structure. Proportion of major workers was most strongly associated with regional climate, consistent with TPH predictions. Colonies invested less in costly major workers in colder, drier environments (figure 3; electronic supplementary material, table S4). While colony size did not vary with climate, it contributed to predicting other colony-level traits. Mean major worker body size was positively related to colony size (electronic supplementary material, figure S1, table S4). Worker body size variance increases with colony size, whereas head size variance was negatively associated with climate and positively with colony size. Head-to-body ratio variance showed a weak relationship with regional climate only.

**Figure 3.**
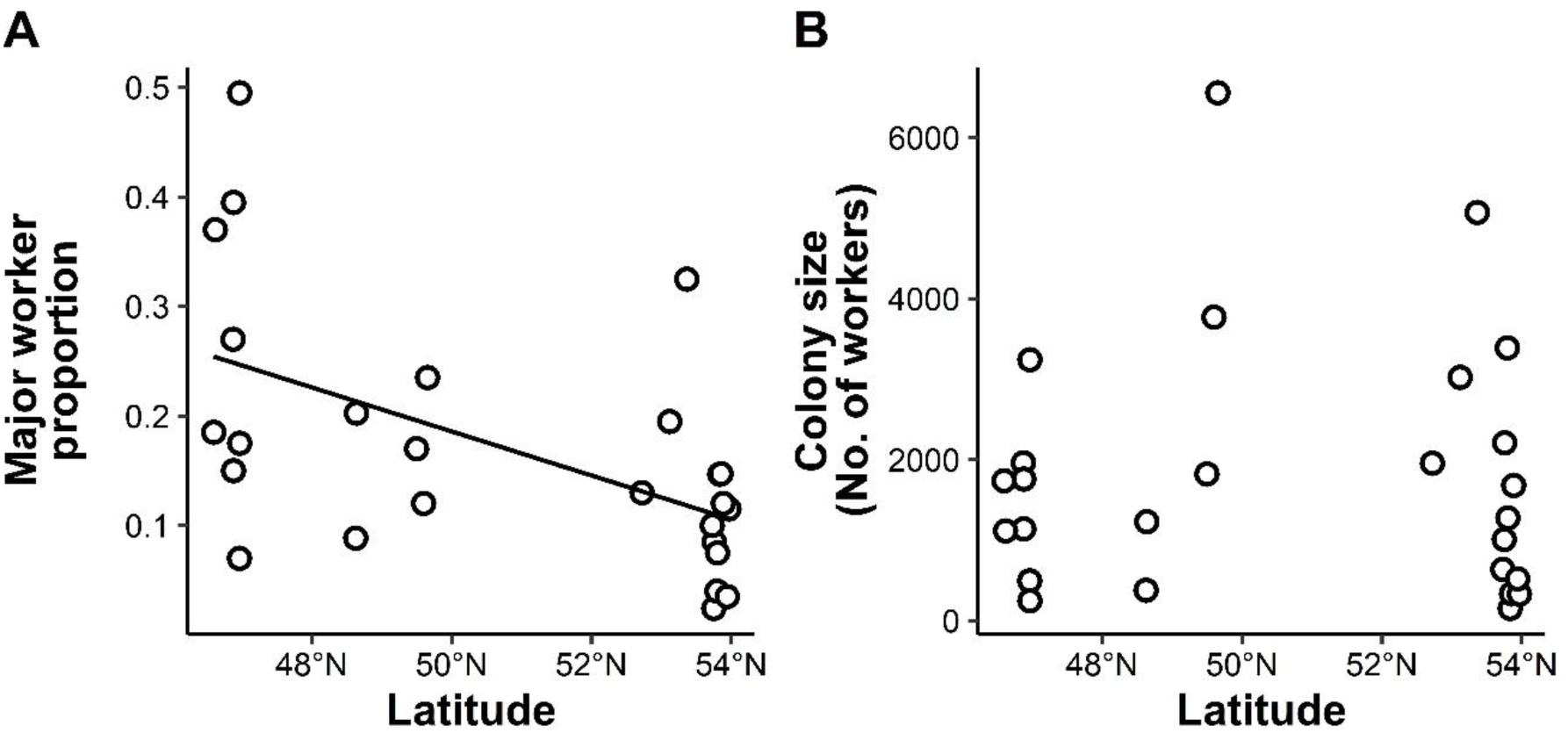
The (A) proportion of major workers and the (B) number of workers in *Camponotus herculeanus* colonies (*n* = 26) in relation to latitude.

**Figure 4.**
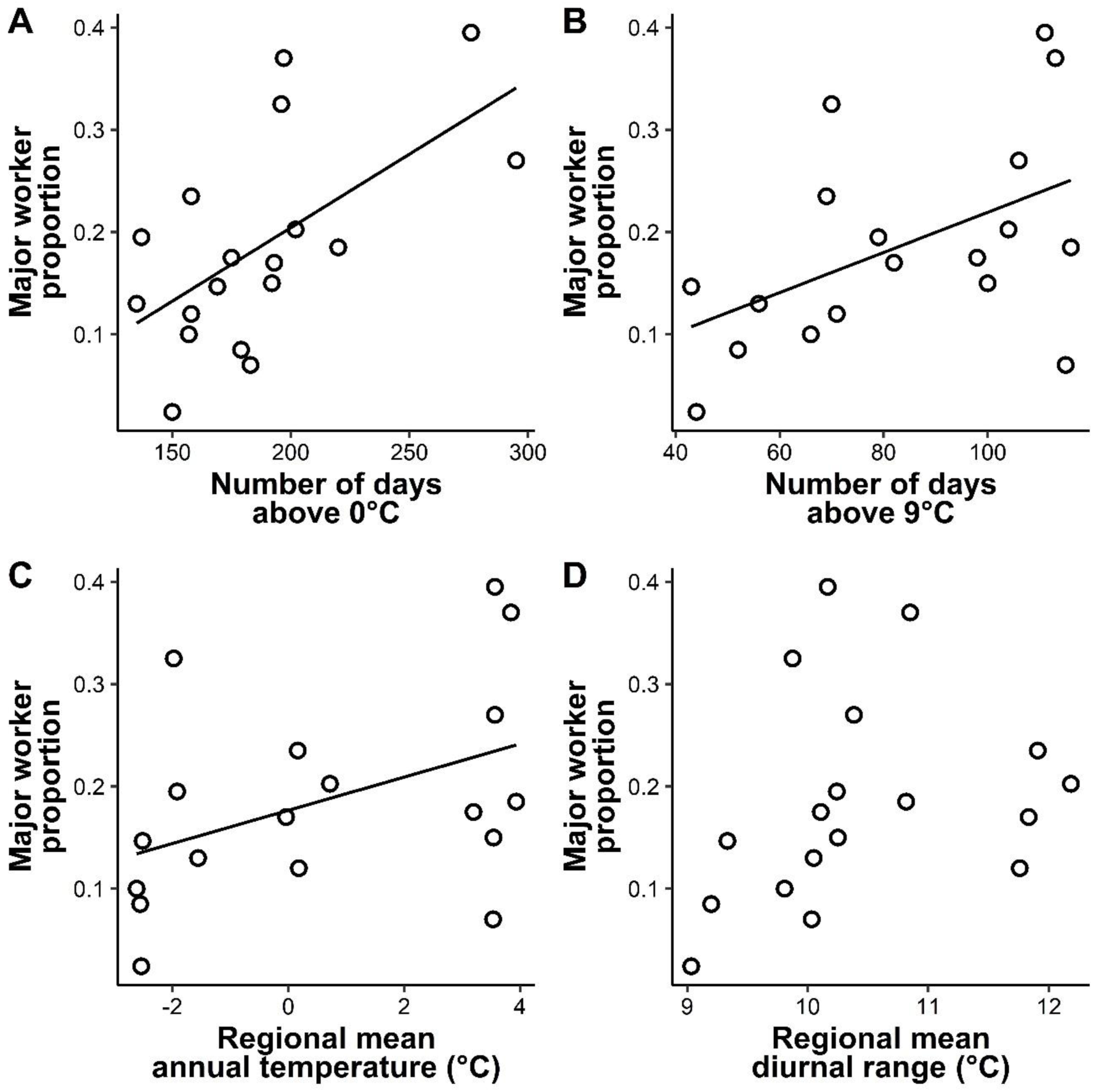
The proportion of major workers in relation to micro- and macro-climate for (*n* = 18) *Camponotus herculeanus* nest site locations. Micro-climatic variables (A & B) were obtained from Thermochron iButton Devices positioned adjacent to each nest. (A) colony activity is estimated as the number of days above 0°C and (B) larval development is estimated as the number of days above 9°C. These temperature measures were obtained between August 21st, 2021, and June 30th, 2022. Macro-climatic variables (C & D) were (C) regional mean annual temperature and (D) regional mean diurnal range for the years 1970 to 2000 [92].

We further tested which temperature factor best predicted the major worker proportion using separate GAMs. The best fit model included the temperature threshold for colony activity, measured as the number of days above 0°, and colony size (electronic supplementary material, table S5). Both predictors showed significant relationships with the proportion of major workers (figure 4a). The second-best model included the temperature threshold for larval development, measured as the number of days above 9°C and colony size (figure 4b; electronic supplementary material, table S5).

Finally, we explored whether colony size is influenced by environmental factors other than climate. Across 26 colonies, colony size ranged from 151 to 6,557 workers (mean ± SE = 1,809 ± 299). Most colonies nested in black spruce (*Picea mariana* (Mill.) BSP; ∼78%), with others found in balsam fir (*Abies balsamea* (L.) Mill.; ∼7%), eastern white cedar (*Thuja occidentalis* L.; ∼7%), eastern white pine (*Pinus strobus* L.; ∼4%), and tamarack (*Larix laricina* (Du Roi) K. Koch; ∼4%). Colony size was positively related to nesting tree diameter (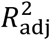 = 0.29, df = 23, *F* = 10.86, *p* = 0.003) and negatively related to the presence of strong competitors, *Formica* spp. (*t* = 2.77, df = 19.66, *p* = 0.01; electronic supplementary material, figure S3). No significant effects were observed for the presence of *Myrmica* (*t* = -1.16, df = 18.26, *p* = 0.26; electronic supplementary material, figure S3) or *Leptothorax* spp. (*t* = 1.01, df = 23.52, *p* = 0.32; electronic supplementary material, figure S3). For additional results on competition estimation, see electronic supplementary material, S5.

## Discussion

Our results support the TPH, which provides a general mechanism by which (super)organisms adapt to extremely stressful environments. We show that eusocial colonies of a geographically widespread ant species consistently reduce their investment in costly major workers when confronted with short growing seasons and most likely, nutritional shortage. Moreover, colonies converged in caste structure after one generation in laboratory settings, under common environmental conditions, which shows that caste structure is a plastic colony-level trait. Overall, our study lends support to the hypothesis that organisms experiencing nutritional shortage in early life stages can plastically reprogram development to rapidly adapt their phenotype to ambient condition.

### The thrifty phenotype as an adaptive response to energetic constraints

The reduction in major worker proportion we observed in northern latitudes, where the growing season is short and food resource are scarce, aligns with the idea that those stressful conditions experienced early during larval development favor energy-saving strategies. Moreover, the absence of relationship between proportion of majors and colony size indicates a reallocation of resources within colonies rather than a general reduction in whole-colony productivity. Given that large colonies have lower per-capita metabolic demand than small colonies [61,62], and that major workers consume more energy per unit of biomass than minor workers [63], energetic shortage in northern colonies could lead to a strategy where colonies maintain their size but reduce the production of major workers. Producing fewer major workers may therefore help colonies maintain overall size while conserving energy, much like individual organisms reduce allocation to costly organs under nutritional stress, which is consistent with patterns observed in vertebrate models of the thrifty phenotype [38,39].

Our study provides insight into the environmental factors that could cause a restructuring of the worker caste. We expected that micro-climate could limit the production of majors by either limiting the amount or quality of resources available to larvae [41,64,65] or limiting the time for development [66,67]. The number of days with temperatures above 0°C was the best predictor of geographic variation in caste structure, indicating that constraints on worker activity, rather than duration of larval development per se, most likely limit the production of energetically costly majors. Thermal constraints on worker activity could limit the amount of time that workers devote to foraging and nursing larvae yearly, possibly reducing the amount or quality of food received by larvae [68–73]. In sum, these results further support the role of nutrient shortage in shaping whole-colony phenotype, possibly via developmental plasticity.

The TPH posits that reduced investment in costly structures is plastic and occurs during development. Consistent with this idea, colonies collected from northern latitudes where foraging and nursing activity of ant workers is limited by the short growing season, produced more majors when reared under warmer and stable controlled conditions with abundant resources, whereas southern colonies produced fewer majors under the same conditions. After one generation under controlled conditions, colonies from different climatic origins converged in caste structure, demonstrating that the reduced investment in major workers observed in the field is environmentally induced rather than genetically fixed. In contrast, body size remained unchanged after rearing, supporting the idea that it is developmentally canalized or under stronger genetic control, and thus decoupled from variation in major worker investment [74]. These results contrast with previous work where population-level differences in worker caste structure across climatic gradients were largely genetically determined [50], possibly due to the much less stressful conditions experienced by ants in this study conducted at more southern latitudes. In sum, our common garden experiment supports the view that organisms can rapidly change their phenotype to match ambient conditions via plastic developmental changes rather than microevolution. Taken together, studies on colony-level adaptation suggests that the balance between plastic and genetic mechanisms regulating worker polymorphism may be evolutionary conserved at the genus level [75–79], with *Camponotus* generally favoring plastic responses to environmental variation [58,64,80–82]. More broadly, our results highlight that adaptive demography in ants can rely heavily on plasticity when environmental predictability is low, and that the relative contribution of plastic versus genetically fixed mechanisms may differ among taxa and environmental contexts.

### The thrifty phenotype as a conserved evolutionary response to environmental stress

Environmentally induced worker caste adjustment could reflect a general and evolutionary conserved mechanism by which organisms rapidly adapt to environmental stress via developmental plasticity [83]. It has been suggested that extremely stressful environments could shape phenotype owing to environmentally sensitive developmental windows during which fine-tuning of developmental pathways to cope with current or anticipated conditions can occur [84]. This view is consistent with evolutionary conserved developmental windows documented in vertebrates and other taxa, where environmental cues, like temperature or nutrient availability, modify developmental trajectories [85,86]. In social insects, and ants in particular, caste differentiation is known to be influenced by temperature, nutrition and hormonal switches [41,57], suggesting that similar conserved developmental mechanisms could underlie the plastic response we observed.

While most studies of the TPH have focused on humans, parallels exist in other model organisms, including rodents. For example, experimental work in mice shows that early-life nutritional restriction results in altered metabolic and growth profiles consistent with a thrifty strategy [60]. In rats, maternal undernutrition induces a clearly defined “thrifty liver programme” characterized by reduced relative liver growth, inflammatory and transcriptomic changes interpreted explicitly within the thrifty phenotype framework [59]. These examples, alongside our findings in natural ant colonies, suggest that the thrifty phenotype may reflect a broad and evolutionarily conserved response to environmental unpredictability across both solitary and superorganisms.

### The thrifty phenotype as an adaptation to unpredictable environments

The thrifty colony phenotype could therefore not only be an energy-saving plastic adaptation, but also as a finely tuned developmental strategy to cope with unpredictable resource availability [4]. The thrifty colony phenotype in *C. herculeanus* illustrates how colonies can adjust caste allocation in a time-sensitive, predictive manner (Figure 6).

**Figure 5.**
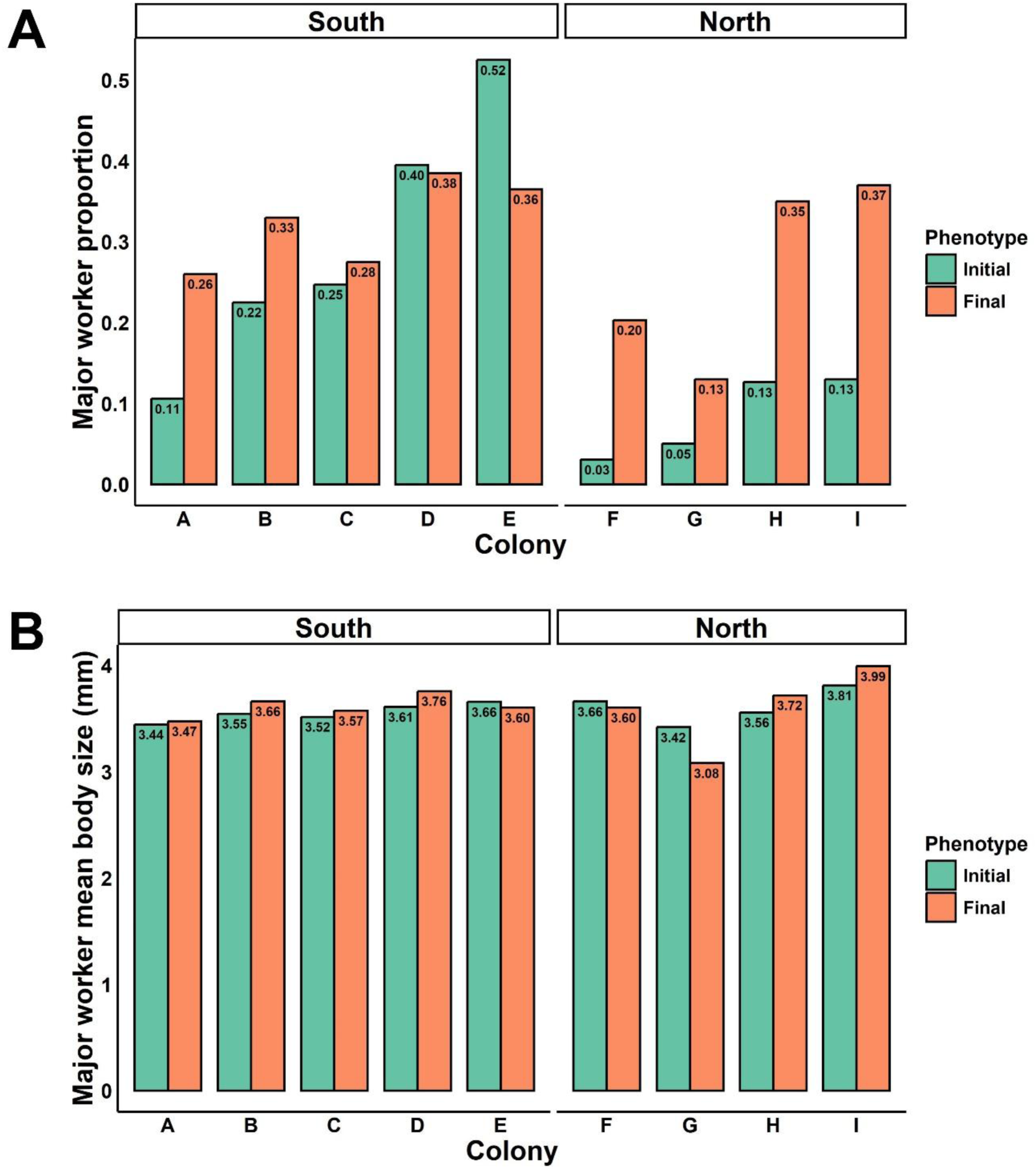
Results from the common garden experiment. Initial (green) and final (orange) (A) proportion of major workers and their average body size (B) in (*n* = 9) *Camponotus herculeanus* colonies collected from the southern (A-E) and northern (F-I) parts of the environmental gradient.

**Figure 6.**
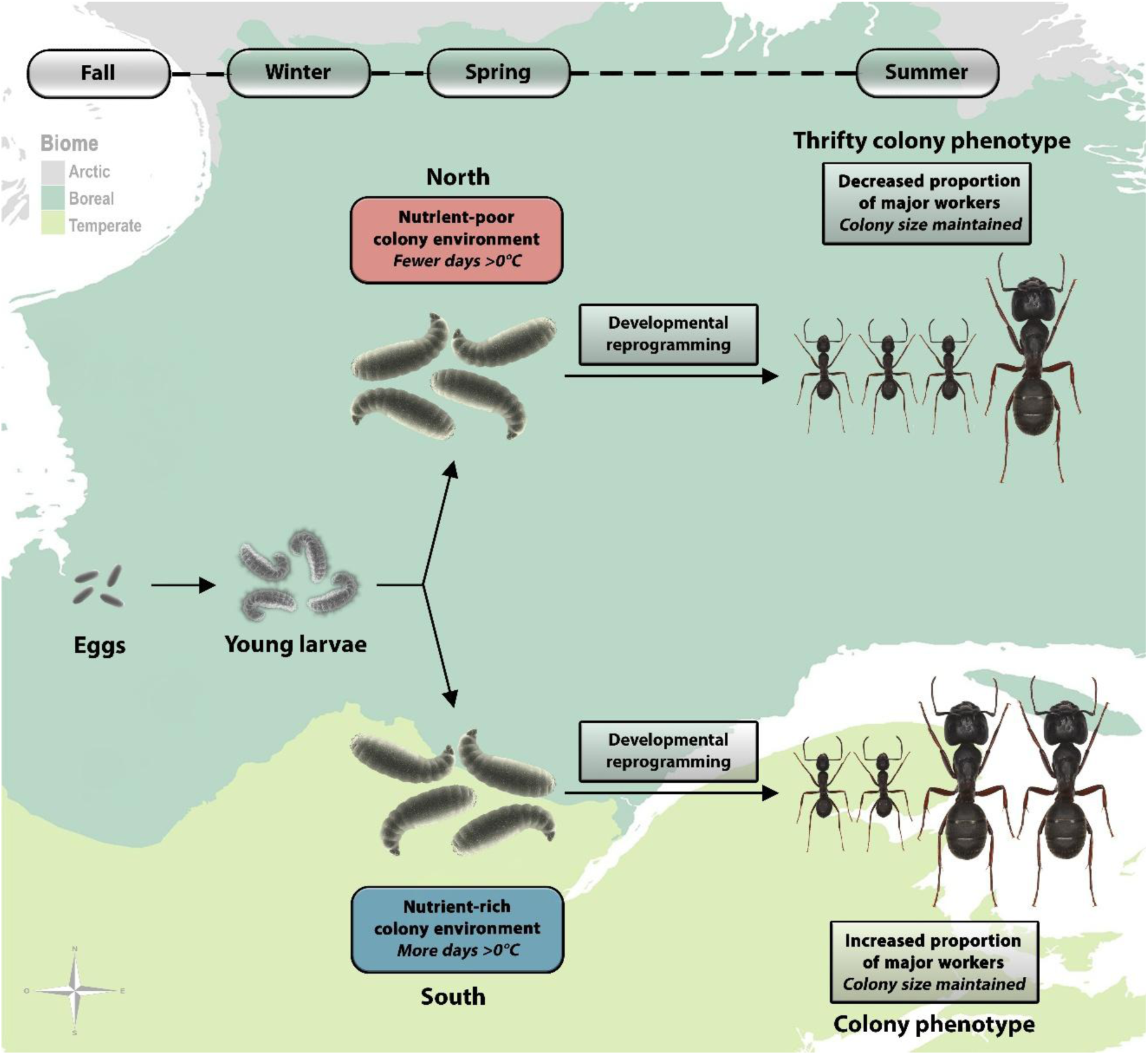
The thrifty phenotype as an adaptation to unpredictable environments in *Camponotus herculeanus*. The figure illustrates colony-level developmental plasticity leading to thrifty phenotype in ant colonies and its nutritional context-dependent consequences on colony survival. Young (early-instar) larvae develop from eggs during fall, enter diapause and receive environmental cues in spring that influence caste determination, potentially mediated by colony-level nutritional conditions. In northern colonies, which experience nutrient-poor environments during the spring, colonies favor the production of cost-effective minor workers over costly major workers (i.e., thrifty phenotype), which may buffer colonies against resource scarcity during nutrient-poor summers, while maintaining colony size. In southern colonies, larvae benefit from nutrient-rich springs, leading to developmental reprogramming and production of more major workers, which should enhance colony fitness during nutrient-rich summers while maintaining colony size. Solid arrows in the figure indicate putative causal pathways from environmental cues to subcaste outcomes.

Bipotent larvae enter diapause during fall, then overwinter, and resume development in spring, when early-spring nutritional conditions possibly provide the critical cue for caste reprogramming. This cue possibly determines whether larvae develop into costly major workers or less expensive minors, effectively predicting the resource environment for the coming active season [58,82]. In populations inhabiting northern latitudes, persistent nutrient scarcity at this early life stage could reprogram larval development to produce fewer majors, generating a thrifty colony phenotype well-suited to short, resource-limited summers. However, if spring cues indicate scarcity while nutritional resources turn out to be abundant during summer, colonies may be maladapted. That is, their low proportion of majors could limit their ability to exploit abundant resources and could even increase mortality due to competition with other nearby colonies. Conversely, when spring cues signal abundant resources, larvae develop more majors, which is advantageous if conditions remain favorable but potentially costly to maintain if resources decline later in the season. The thrifty colony phenotype could therefore be a predictive developmental adjustment, with fitness payoffs dependant on the reliability of environmental cues across life stages.

Environmental cue–response systems are predicted by models of predictive adaptive plasticity and seasonal polyphenism, in which developmental trajectories are adjusted based on early-life environmental signals to optimize performance in anticipated conditions [87]. When environmental predictability is low, these same systems can produce mismatch costs, a form of developmental bet-hedging that may help explain variation in colony success at the northern range edge of the species, where environmental conditions are expected to be extremely stressful [88].

### Building on previous theory about colony-level adaptation

Our findings further shed new light on the validity of one of the key hypotheses proposed to explain geographic variation in body and colony size in eusocial organisms. The *starvation resistance hypothesis* predicts that colonies experiencing long winters should rely on larger workers and/or larger colonies to store fat and nutrients, thereby buffering winter mortality [89,90]. In our study, however, neither body nor colony size varied with climate. Only caste structure, namely the proportion of major workers produced by colonies, co-varied with climatic conditions. This contrasts with Yang et al. (2004), who found that in colonies of *Pheidole morrisii* Forel, 1886, the proportion of majors declined in more seasonal climates, but that the average size of majors increased [50]; in those populations, majors in seasonal climates often functioned as repletes for fat storage [91]. Together, these studies suggest that both the *starvation resistance hypothesis*- and the TPH-related patterns can occur depending on the species and ecological context, whereas our results are more consistent with the TPH.

## Conclusions

Our study presents the results of the first empirical test of the TPH performed in natural systems outside humans. The findings of developmentally plastic thrifty phenotypes in superorganisms inhabiting northern latitudes could reflect both energetic constraints and adaptive, predictive responses to environmental stress, similar to what is observed in humans, supports the view that the thrifty phenotype strategy is evolutionary conserved between insects and humans [23]. The potentially conserved developmental plasticity observed here in a superorganism may offer a powerful model to study how stress-responsive phenotypes evolve in complex, long-lived organisms, including humans [28].

## Methods

### Study sites

The study took place in the province of Quebec, Canada (figure 2). Study sites spanned from temperate deciduous forest of southern to the northern limit of the boreal forest (aka, tree line). The southern sites were in the Mauricie (46.6°N, -72.5°W) and National Capital (46.8°N, -71.5°W) regions whereas the northern sites were above 52°N in Northern Quebec region. Along this environmental gradient, mean annual temperature ranges from 3.93°C to -2.79°C, minimum annual temperature of coldest month ranges from -18.10°C to -28.70°C, and annual precipitation ranges from 1383 mm to 572 mm [92]. The growing season are approximately 140 to 150 days in the south and 100 to 110 days in the north [93].

### Study organism

We used *C. herculeanus* (figure 1) as a study organism because it is the most broadly distributed, polymorphic eusocial species in eastern Canada for which it is possible to collect entire colonies. *C. herculeanus* is widely distributed throughout the Holarctic realm [94]. In the Nearctic region, entire colonies of this species can often be found inside a single tree, stump or log and frequently occur in peat bogs [95]. It builds its nest by digging chambers inside dead or live wood. In addition, since peat bogs are frequently flooded, the colonies rarely ramify into very complex tunnel systems or satellite nests in these habitats, thus allowing the complete collecting of the colony. *C. herculeanus* is a monogynous (single egg-laying queen), slow-growing species and the most cold-tolerant ant species in the world [95,96]. Based on Wilson’s classification [56], *C. herculeanus* exhibits continuous worker polymorphism, specifically triphasic allometry, producing clear minor and major worker subcastes (in line with recent evidence in *Camponotus* [58]). Individuals falling between these endpoints (intermediate workers or medias) are best interpreted as vestigial products of developmental threshold slack rather than a distinct subcaste. For details on identification of the major worker subcaste, see electronic supplementary material, S1.

### Colony collecting for observational study

For the observational study aiming to document several colony-level traits, we collected 26 colonies in peat bogs along the environmental gradient. We only collected colonies including at least 150 workers. Colonies with less that 150 workers were not collected because they are considered incipient and rarely contain major workers [97]. When excavating the colony, we collected all workers, broods and alates we observed. Colonies were collected at least 1km apart. We collected the northern colonies (∼52°N to ∼54°N) from mid-July to early August, and southern colonies (∼46°N to ∼52°N) in both mid-July and early August of summer 2021. These dates were selected as to not bias our estimation of colony traits because of winter mortality. They correspond with periods when the yearly generation was hatched in the colonies [98,99]. For more details on colony collecting, see electronic supplementary material, S2.

### Colony collecting for common garden experiment

To test whether the variation in the caste structure along the gradient is due to microevolution or developmental plasticity, we collected nine entire colonies along the same gradient and using the same methodology as in the observational study. In 2022, we collected five colonies in the southern sites (September 16^th^ to 25^th^), and four in the northern sites (August 4^th^ to 9^th^). Colony size was estimated visually by counting the number of occupied plaster nests as to not stress or hurt workers. The average colony size was similar (1,667 ±333 (*SE*) workers) to the average size of all colonies we ever collected along this gradient (1,772 ±299 (*SE*) workers) (includes unpublished observations).

### Estimating regional climate

To estimate variation in regional climate, we extracted several georeferenced bioclimatic variables that could possibly influence worker polymorphism. Bioclimatic variables were extracted for each site where a colony was collected using WorldClim data at 1 km^2^ spatial resolution, which included mean annual temperature, mean diurnal range, isothermality, temperature seasonality, maximum temperature of warmest month, minimum temperature of coldest month, temperature annual range, and annual precipitation [92,100]. We performed a principal component analysis (PCA) on the eight bioclimatic variables listed above to reduce the dimensionality of the data and collinearity [101]. We extracted the first axis, which captured 76% of the variance (electronic supplementary material, table S1).

### Estimating local conditions

The size of nesting spaces can impose limits on colony size in carpenter ants and possibly the degree of worker polymorphism [102,103]. We therefore measured nesting tree diameter at the base of the tree trunk. We also estimated canopy openness using a spherical densiometer (Forest Densiometers, Model-C, Bartlesville, Oklahoma, USA) as it relates to the amount of light reaching the nest, thereby altering its temperature and the amount of energy and food resources available. In turn, temperature can influence larval growth rate [104] and whether some larvae might pupate or enter diapause [105], which could influence worker polymorphism and colony size in *C. herculeanus* colonies.

### Estimating local temperature

To estimate the temperature regime experienced by colonies, we used Thermocron iButton DS1921G-F5# integrated thermometer and iButton data-logger (Maxim Integrated, San Jose, California, USA). They were deployed at 18 of the 26 colonies collected in 2021 due to resource and logistical limitations. Temperature was measured every 255 minutes (maximum length for the data-logger model) for 12 months. We then estimated the number of days during which brood development is possible (above 9°C) and the number of days during which worker activity is possible (above 0°C) (for more details, see electronic supplementary material, S3).

### Estimating the intensity of competition

The presence of competitive ant species near *C. herculeanus* colonies could influence major worker production, as major workers can have a role for defense of the colony [98]. We assessed the presence of potential interspecific ant competitors by active visual observation and opening wood debris within a radius of ten meters around each colony for 15 minutes between 10:00AM and 5:00PM on days without precipitations. We sampled workers of all species found with a mouth aspirator or manually. Species identifications were confirmed under a microscope by a trained ant taxonomist.

### Morphometric measurements

To estimate colony-level traits, we took morphometric measurements on 200 randomly sampled workers in each colony, except for one colony comprising a total of 151 workers, for which we measured all the workers. We froze collected colonies within five days of collecting to avoid influencing the proportion of major workers or any other colony-level traits. To avoid selection bias, we created all the subsets on the same day using the same method by mixing all the workers of each colony together in a plastic container with ethanol for two minutes and randomly collecting 200 workers using forceps.

We took photos of morphological traits using a digital microscope (Brand: Koolertron, Model: AD106S). We digitalized the dorsal and lateral views of each specimen and obtained morphometric measurements using an image processing software (ImageJ version 1.53e). For each worker, we measured head width (maximum width of head including eyes) and body size (Weber’s length, i.e. maximum diagonal length of mesosoma) (electronic supplementary material, figure S4).

### Measuring colony-level traits

We first estimated colony size by counting each collected worker using forceps and a tally counter. We then took morphometric measurements to quantify five different colony-level traits describing worker polymorphism, including the proportion of major workers, mean major worker body size, worker body size variance, worker head size variance and worker head-to-body ratio variance. We quantified five different traits related to worker polymorphism because each of them related to different functions for colony adaptation (table 1). We calculated the mean size of major workers as the average of body size measurements (i.e., Weber’s length) of major workers in the entire colony. We calculated the variance of worker body size (i.e. Weber’s length), head size, and the ratio between head width and worker body size for each of the sampled ant colonies. We used variance because it is the parameter that best captured the variation of each trait between the colonies (electronic supplementary material, table S3) and represents how large the variation of worker morphology is in comparison to the other workers of a same colony.

**Table 1.** Rationale for using different ant colony-level traits of the worker polymorphism.

| Colony trait | Functions for the colony |
| --- | --- |
| Proportion of major workers | **Defense of the colony** [50,89,112–117] |
|  | **Food storage** [89,91] |
| Major worker mean body size | **Thermal tolerance** [118–123] |
|  | **Desiccation** [124,125] |
| Worker head and body size variance | **Dietary composition** [126–129] |
|  | **Division of labor** [56,57,130] |
| Worker head-to-body ratio variance | **Habitat complexity** [131–134] |
|  | **Division of labor** [135–138] |

### Common garden experiment

We ran the common garden experiment from August 10^th^, 2022, to June 21^st^, 2023. For detailed information on the experimental conditions, see electronic supplementary material, S4. We focused this experiment on the proportion of major workers as a response variable because it was the only colony-level trait that varied independently from colony size and was predicted by an environmental variable. To obtain a representative estimation of the proportions of major workers without harming the colonies, we randomly selected 200 workers in each colony upon collection in nature. To do so, for each colony, we placed all adult workers in a same plastic box and randomly sampled them using forceps while not looking directly at the sample and shaking the box every 30 seconds to avoid worker clustering. This procedure ensured that our subsample was not biased toward a particular subcaste that tend to cluster or that might be inadvertently chosen when sampling visually [106]. Then, to compare the colony phenotypes, we waited until brood develops into adulthood after diapause in a common rearing environment and randomly resampled 200 workers using the same method after one larval generation.

### Statistical analyses

#### Environmental predictors of colony-level traits

To select the variables to be used in a multivariate normal additive model relating worker polymorphism to environmental factors, we ran separate generalized additive models (GAMs) for each of the five colony-level traits [107]. We included the predictor variables of regional climate, local climate, nesting tree diameter, colony size and competition (i.e., presence of competitor). After fitting a separate GAM for each trait, only the significant predictor variables were kept for the multivariate normal GAM.

The multivariate normal GAM fits a separate regression for each response variable, as a covariance matrix estimating the covariance between response residuals. The output includes a separate linear predictor for the mean for each response and the covariance matrix. The conditional mean in a GAM is estimated as a linear combination of nonlinear basis functions, that form a smooth curve. GAMs are adequate for selecting variables because p-values can be used for testing each variable for equality to zero for deciding on which candidate variables to be removed from the model.

Multivariate normal GAMs can be used to assess the relationship between many variables and are robust to autocorrelation. Using only the selected variables, we fitted a single multivariate GAM [108]. To fit this multivariate model, we used the five colony-level traits as response variables representative of ant worker polymorphism (table 1). We used “colony size” and “regional climate” as fixed effect predictor variables for each response because they were the significant predictors in the previous approach using separate GAMs for each colony-level trait. We used linear predictors instead of smoothing terms based on the model linear relationships between response and predictors residuals. Finally, we assessed the relationship between each colony trait and the predictors, and we plotted the partial effects as a visual aid for the direction of the relationships (electronic supplementary material, figure S6). We fitted the model using the *gam* function in the *mgcv* package [109].

To test which temperature factor best predict the proportion of major workers, we ran four separate GAMs with the proportion of major workers and colony size as response variables and the number of days above 0 °C, the number of days above 9°C, the regional mean annual temperature (°C) and the regional mean diurnal range (°C) as predictors. Then, we determined which was the best-fitting model based on the lowest corrected Akaike’s information criteria (AICc) value, adjusted for small sample size. We calculated AICc values using the *AICc* function in the *AICcmodavg* package [110].

#### Environmental predictors of colony size

To assess the relationship between colony size and the environmental variables (predictors) we used a linear model using the *lm* function. We performed Welch two-sample t tests to test whether colony size differed in the presence of certain ant genera using the t.test function.

#### Plasticity of caste structure

During the common garden experiment, to test whether the proportion of major workers or mean major worker body size significantly changed from the moment of collection to after one larval generation in a common rearing environment, we performed Welch two-sample t tests for each group (North and South) using the *t.test* function. We also tested whether the proportion of major workers in both groups were significantly different from each other using the same method. All statistical analyzes were performed in R version 4.2.0 [111].

## Acknowledgements

We are especially grateful to D. Ouellette (McGill University) for his assistance during fieldwork and for mounting specimens in figure 1. We also thank E.J. Pedersen, E. Despland, and G. Muñoz (Concordia University) for their advice on analytical approaches, as well as P.-M. Brousseau (Concordia University) and members of the Abouheif Lab (McGill University) for their helpful comments on the manuscript. We also thank K. Marshall (University of British Columbia) and R. Khelifa (Concordia University) for valuable discussions. We acknowledge the Centre for Northern Studies for providing access to research infrastructure and the McGill University Phytotron facility for support of the common garden experiment.

