## Supplementary material for "Thrifty phenotypes in ants: Extending a human developmental hypothesis to a superorganism"

### Supplementary Notes

*Supplementary Note 1: Identification of major workers*

Majors are larger than minors or medias with a disproportionally large head relative to their body size (figure 1; electronic supplementary material, figure S5). It is therefore easy to quantitatively differentiate the castes, which is useful for studying variation in caste structure [1]. As the workers exhibit a continuous size variation, the sub-castes cannot easily be distinguished visually. The inflection points of the allometric growth curves can be used to classify *C. herculeanus* workers into minor, media, and major's sub-castes. The allometric growth curve of *C. herculeanus* is a third-degree polynomial formed when plotting head size against body size (electronic supplementary material, figure S5). The two inflection points at which the derivative of the curve is equal to zero represent the separation between the three worker sub-castes, allowing to classify them [2]. When plotting head size against body size using the measurements of the workers of a same colony, minors are those with a body size smaller than the body size at the first inflection point, majors are those with a body size larger than the body size at the second inflection point, and medias are those in between those inflection points. In this study, we only calculated the proportion of major workers using the number of workers as those with a body size larger than the body size at the second inflection point (electronic supplementary material, figure S5).

*Supplementary Note 2: Colony collecting for observational study*

Based on our observations, *C. herculeanus* is commonly found on the edge of peat bogs. For the observational study, we first selected suitable sites using Google Maps. Most of the collecting sites were located at least 100 meters from roads, trails, overhead power lines or any form of human disturbance, to ensure each colony is in natural conditions. When the species was present in a selected site, we collected at the edge of peat bogs for sites to be as comparable as possible.

We collected the colonies during summer 2021, from mid-July to mid-August, when they were active and re-established after swarming, winter mortality and spring brood hatching to not bias the colony traits [3,4]. Winter mortality is caused in part by predation by several species of woodpeckers and may be important enough to influence colony sub-caste ratios, with *C. herculeanus* being a major part of their winter diet [5–7]. Winter mortality in workers might also be caused in part by cold temperatures, as other ant species can use workers to buffer the queen against cold temperatures [8,9].

To collect entire colonies, we selected tree trunks that we could open completely without danger of injury due to the top part falling and contained the desired species to collect entire colonies. *C. herculeanus* is known to prefer softwood, especially *Abies, Picea* and *Thuja* species to build their nests [6,10–12]. We opened trunks with a saw and an axe on a tarpaulin on the ground and we directly deposited the debris containing the most individuals in ventilated plastic containers identified with the unique code of the colony. We collected other moving individuals manually. Due to the high summer temperatures during collecting, ants can die quickly when they are numerous and active in a closed container. Therefore, within 30 minutes after collecting, we placed the colony containers inside a cooler, and then in the refrigerator (4°C) when returning from the field to prevent them from becoming too active. We collected the individuals manually or with forceps from the debris in open plastic bins. Talc powder mixed with anhydrous ethanol was applied to the walls of the containers so that the ants did not escape during the transfer. Then, we isolated the ants to only have the individuals that we euthanized and kept in 95% ethanol until we took their morphometric measurements. We confirmed *C. herculeanus* identification under a microscope using the criteria mentioned in Ellison *et al.* (2012).

The collecting dates were important to ensure that the number of major workers found during collecting reflects the maximum that can be produced by the colony during the season. Therefore, we set collecting dates after the first brood development and based on differences in the onset and length of the growing season.  In fact, *C. herculeanus* first brood development is independent from temperature and the second brood usually mostly enters diapause [13]. The main period of larval growth at latitude 46 ° and above in Eastern Canada are in June and early July and the remaining larvae often spend the winter in the nest as the life cycle can cover two years [6,14]. Since freezing temperatures are still a possibility until mid-June in the northern part of the environmental gradient (above 49° N), collecting was not conducted until mid-July to make sure that a full month has elapsed since the last frost and that worker brood was hatched.

Mating flights are also an indicator of the beginning of the species activity after diapause and therefore of brood production. In *C. herculeanus*, winged sexual castes spend the winter in the colony and mating flights can occur during spring when temperatures are above 21°C and some snow can be remaining on the ground [6]. Major mating flights of *C. herculeanus* take place between the second week of May and the first week of June in Eastern Canada although some can take place later [6,15–17].

*Supplementary Note 3: Estimating colony-level temperature*

Each Thermochron iButton Device was encapsulated in 50 mL Fisher Scientific polypropylene centrifuge tubes. The tube's lid was sealed with waterproof silicone (ASTM C920 Class 35). The tube was placed in a double-sealed plastic ziploc bag and taped to 8 mm galvanized 26.67 cm steel pegs, five to ten centimeters under the surface of the soil, 10 cm away from each nest. The devices were placed at this position, as most of the adults and brood are located at the base of the nesting tree, down to 15cm underground. Most of the colony also overwinters in the lowermost section of the nest (Sanders 1964). Since the devices only allowed one year of data storage and that the devices were not all placed at the same date across sites, only the data from August 21st, 2021 to June 30th, 2022 was analyzed for the data to be as comparable as possible between sites.

As larger ants generally take longer to develop than smaller ants in other species [18], nests with longer periods of temperature conditions allowing brood development are expected to have a higher proportion of major workers than nests with shorter growing seasons. To test the hypothesis that a lower proportion of major workers represent a potential adaptation to short growing seasons, we calculated the number of days the temperature was above 0°C (temperature threshold allowing colony activity) and above 9°C (temperature threshold allowing larval development) between August 21st, 2021, and June 30th, 2022, at the collecting location of 18 colonies along the environmental gradient. We also compared these same colonies with the regional mean annual temperature and the regional mean diurnal range using the same data extracted from WorldClim for the previous analysis.

The temperature thresholds were chosen by considering several factors which are detailed below, including the minimal temperature for brood development, which is 17°C [13], and the adult activity threshold temperature, which is between 7 and 10°C in Eastern Canada [14]. The temperature inside the nest is also expected to be significantly higher than the external temperature. For instance, in North-east Asia, the temperature inside *C. herculeanus* nests is around 12°C warmer than air temperature in winter [19]. The workers can also move the brood in the part of the tree heated by the sun during the larval development, rising the temperature to speed up the development of workers [14,20].

Sanders (1972) hypothesizes that the temperature inside the nests is controlled by the metabolic heat of the ants after showing that the temperature inside the nest can be raised as much to 16°C above the temperature outside the nest. Consistently with Sanders' hypothesis, the first author observed several thousand workers clustering around the remaining late instar larvae and cocoons inside a rotting stump of 25 cm diameter by 38 cm long in a closed canopy forest during a cold morning on August 1st, 2021, in Eeyou Istchee James Bay, QC (49.5986; 77.3103), and the cluster of ants felt very warm to touch. Eidmann (1942) also observed that workers of *C. herculeanus* begin to cluster at 8°C, and that the cluster increases in density with decreasing temperature, the cluster completing at 5°C. The structure of clusters in *C. herculeanus* is always similar; the queen and brood are in the center, then the workers cluster around them with reproductives irregularly mixed throughout the cluster, and the majors, forming the peripheral layer, are the last to join the cluster, and the first to leave it [21]. While the cluster forms between 8 and 5°C, the majors wonder around the nest, guarding the colony.

*Supplementary Note 4: Common garden experiment conditions*

We kept all the colonies in a Conviron environmental chamber under the same artificial environment setups with controlled temperature and light thermoperiod, humidity, and unlimited food access. We kept the colonies in separate 45.7 × 61.0 × 17.8 cm plastic boxes with two holes of 15 cm diameter in the lid to allow air flow. We covered the holes with glued metal mesh to prevent ants from escaping. We also covered the walls of the boxes with talc powder mixed with anhydrous ethanol to prevent escape. We handmade artificial plaster nests fitted in 100 × 25 mm petri dishes and deposited them at the bottom of the boxes. The number of nests was dependant upon their colonization by the ants. To allow unlimited colony growth, we added nests until there were at least two empty nests. We fed the colonies twice a week with an unlimited amount of Bhatkar-Whitcomb diet [22] and frozen tropical house crickets (*Gryllodes sigillatus* (Walker, 1869)), meaning that there was always food left at each feeding time. We also provided the colonies with unlimited water and sugar-water (250g /L) sources.

We conducted an extensive review of the literature to assess which ambient conditions would be most suitable for the survival of *C. herculeanus* colonies during the common garden experiment. Experiments conducted by Zhigulskaya (1987) show that the optimal temperature for development of the eggs and larvae of *C. herculeanus* is 27°C in the laboratory. It is also reported that some of the second instar larvae do not develop during the second year of their life and must go through a second overwintering period, often during during cold and rainy summers [23,24]. In our experiment, all larvae developed after a single diapause. It is probable that the life-history of *C. herculeanus* is slightly different with the North American populations that we used in our experiment. According to Lopatina and Kipyatkov (1993), the first brood larvae all pupate independently of temperature. However, for the second brood generation, only a few larvae completed their development during their experiments, and exclusively at 25 and 30°C constant temperature and thermoperiods 16/30°C, 14/32°C (cycles of temperature during day and night), but not at a constant temperature of 23°C [13]. Furthermore, 12h thermoperiods were shown to accelerate development, increase queen productivity, as well as brood survival [13]. This made sense with the first author's observations in the field, as *C. herculeanus* often nests in standing dead trees; the nest heats up with the sun during the day, and then gets cooler at night. However, no larvae of the second brood generation pupated in our experiment.

Since the species requires a reasonably long diapause and that we wanted to maximize survival for the experiment, we chose the most optimal experimental conditions at the best of our knowledge, building upon the literature, our observations in the field, as well as our experience in arthropod rearing. We set an 18/27°C 13h thermoperiod with a cycle day and night. We also set infrared lights to be turned on from 5AM to 9PM, which is similar to the month of June's photoperiod in the south of the environmental gradient (5AM to 9PM). The temperature was set to gradually lower, at 8PM (25°C), 9PM (23°C), 10PM (21°C), and 11PM (18°C), and gradually increase at 4AM (21°C), 5AM (23°C), 6AM (25°C), and 7AM (27°C), by increments of 1°C every 30 minutes. Humidity was kept at 40% to prevent mold and mites' infestations, which can negatively affect colony development.

When typical signs showed that all the colonies were ready to overwinter, we began to change the chamber's conditions for beginning the diapause, as larval development cannot pursue unless the workers overwinter [25]. As Hölldobler (1961) described, we observed that independently from the environmental conditions in the chamber, the workers stopped foraging and plugged the entrance of their nests using any material they could find, which here mostly included the Bhatkar-Whitcomb diet. Most of the workers remained inside the nests with the brood and lower levels of activity were observed. We stopped providing the Bhatkar-Whitcomb diet and the crickets, and lowered the temperature by increments of 1°C per day until it reached a constant 4°C. We also gradually reduced the amount of time the red lights were turned on by increments one hour per day until the lights were shut off.

The chamber remained at a constant 4°C with lights off for four months. Typically, endogenous-heterodynamous ant species minimally require one to four months at 3-5°C [26]. *C. herculeanus* specifically requires three to four months of diapause [25]. We refilled water reserves of the nests twice during the diapause to prevent desiccation. After four months, we gradually increased the temperature by increments of 1°C per day until reaching the original conditions prior to diapause. We also increased the duration of red lights by increments of one hour per day until reaching the original conditions.

In the colonies where alates were present, we observed that most of the alates came out of the nests when temperature reached 22°C and that drones began flying inside of the boxes. This is highly consistent with observations by Sanders (1964), who states that *C. herculeanus* nuptial flights normally take place when temperatures are above

21°C. To mimic natural conditions, we manually removed all alates from the boxes on the day that temperature reached 22°C.

According to our observations, all the common rearing environment conditions that were set were unproblematic and no mortality in the colonies' brood was observed at any stage of their stay in the experimental chamber. Upon the end of the larval development, all the pupae hatched successfully into viable adults. Our observations support the idea that the conditions were optimal for all the colonies during the experiment.

*Supplementary Note 5: Potential competitors*

The most common ant genera we found near *C. herculeanus* nests were *Leptothorax*, *Myrmica* and *Formica* (electronic supplementary material, table S6). The main competitors of *C. herculeanus* in peat bogs appear to be *Formica spp.*, such as *Formica neorufibarbis* Emery, 1893 because they are the only relatively large-sized ants occupying the same habitat, nesting in wood logs, and behaving aggressively [27–30]. Other carpenter ant species (e.g., *Camponotus pennsylvanicus* (De Geer, 1773)) and *Camponotus novaeboracensis* (Fitch, 1855) are also potential competitors, but here we did not include them in the analysis because they were not observed near *C. herculeanus* nests. The species *Camponotus nearcticus* Emery, 1893 occurred at one site, but we found no other species of *Camponotus*.

### Supplementary figures

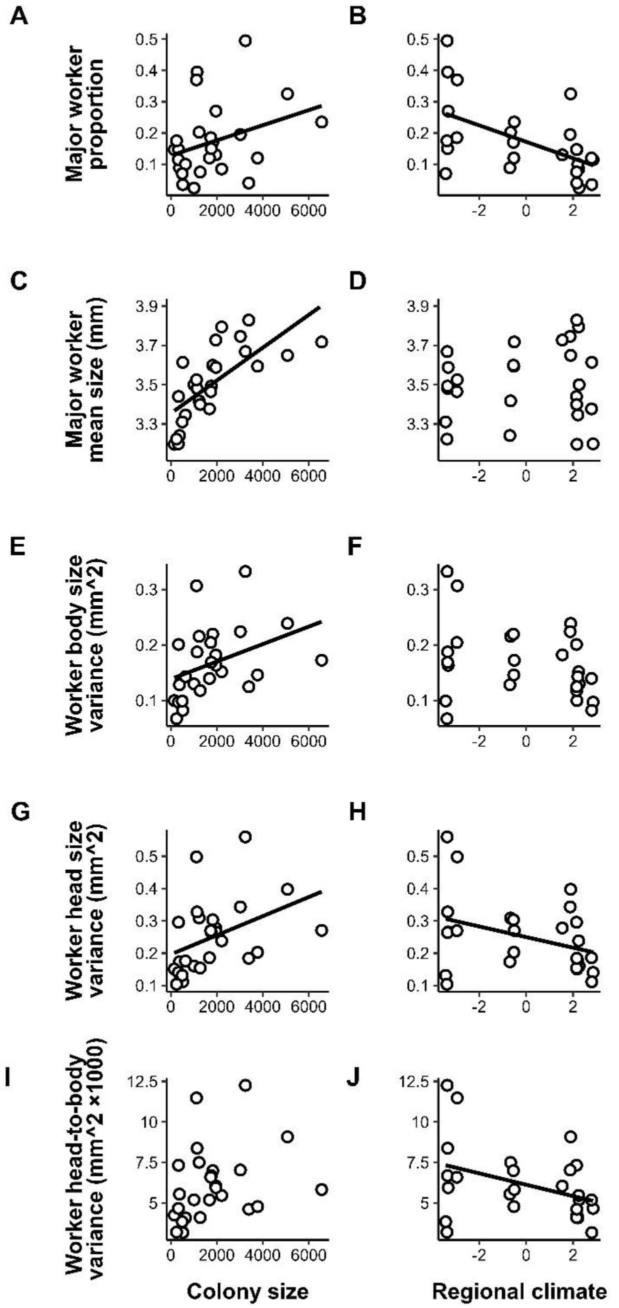


**Supplementary Figure 1.** Colony traits *Camponotus herculeanus* collected in 2021 along an environmental gradient in Quebec, Canada according to colony size (total number of workers) and regional climate (PC1, positive values represent a colder and drier climate, while negative values represent a warmer and more humid climate) ($n$ = 26).

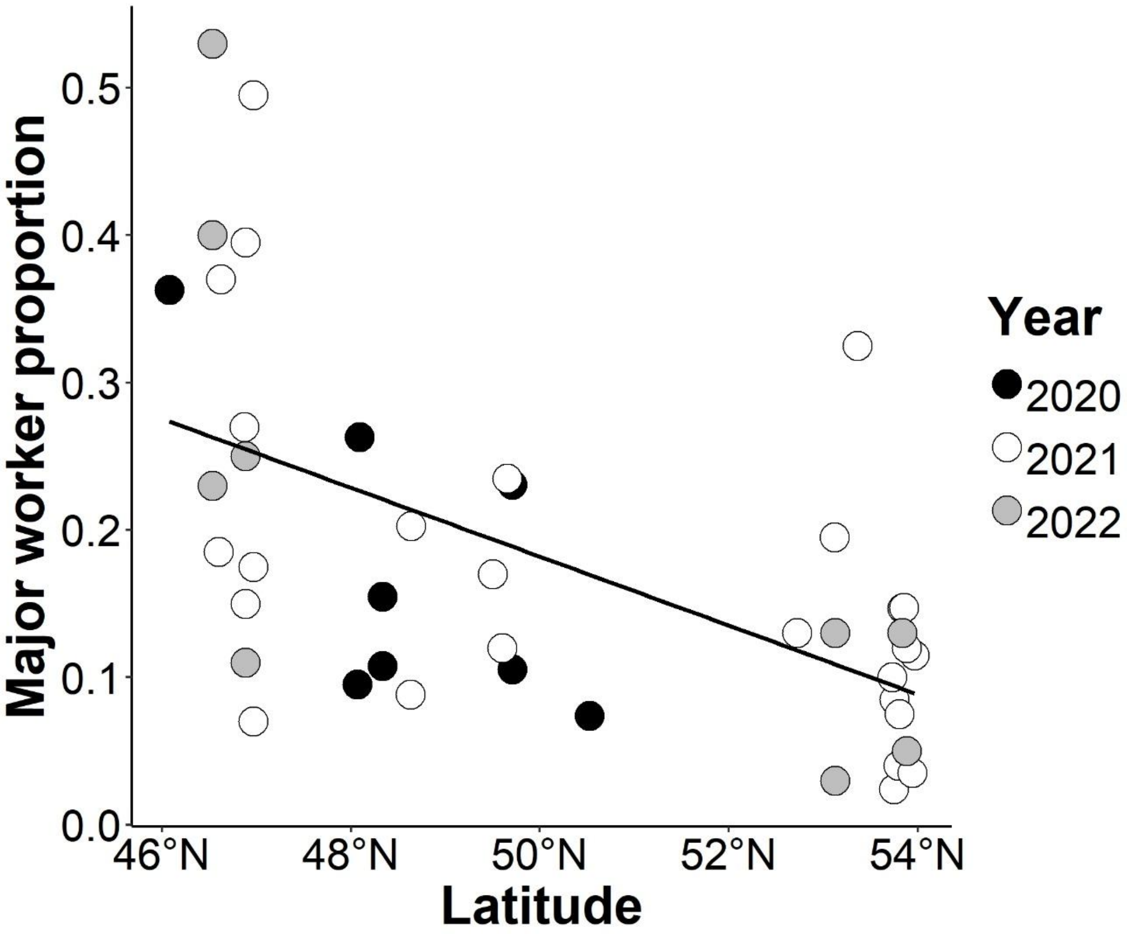


**Supplementary Figure 2.** The proportion of major workers among 44 colonies of *Camponotus herculeanus* along the environmental gradient. Nine colonies were collected in 2020, 26 in 2021 and nine in 2022.

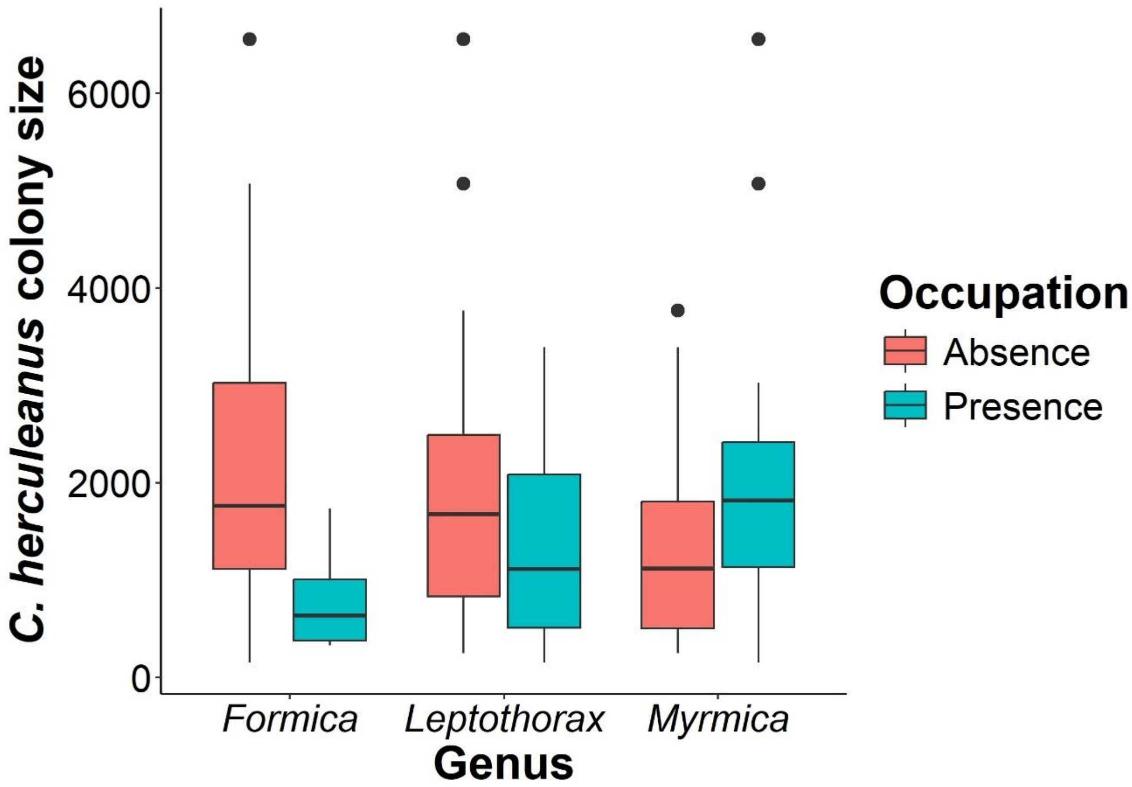


**Supplementary Figure 3.** Size of *Camponotus herculeanus* colonies (number of workers) collected in 2021 along an environmental gradient in Quebec, Canada according to the presence or absence of nests of competitive ant genera in a radius of 10 meters (*n* = 26).

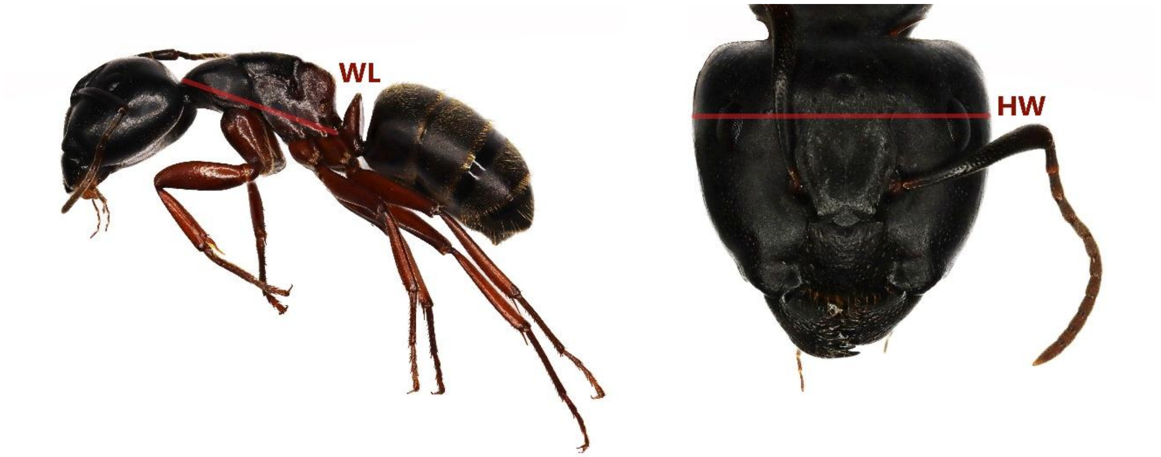


**Supplementary Figure 4.** Weber's length (WL) and head width (HW) of a major worker *Camponotus herculeanus* collected in Quebec, Canada.

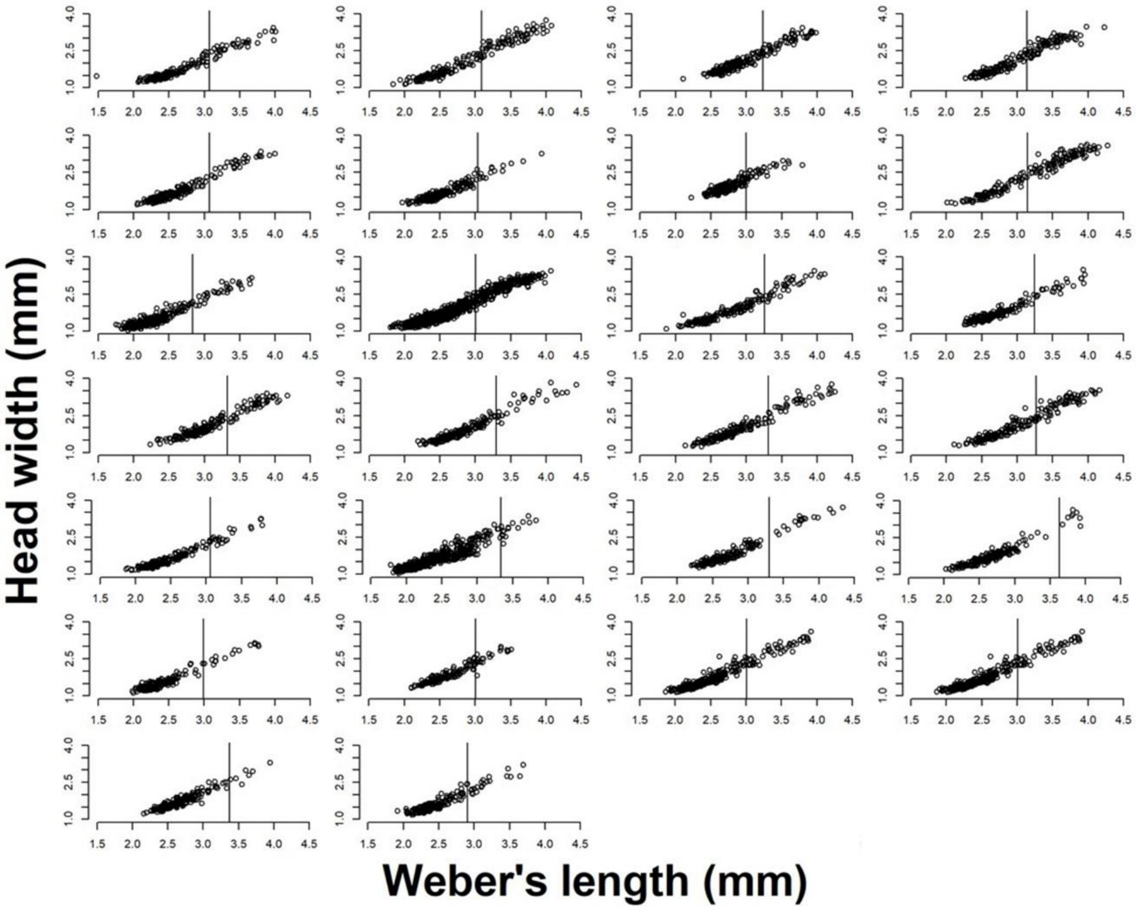


**Supplementary Figure 5**. Second inflection point of the allometric growth curve of 26 colonies of *Camponotus herculeanus* collected in 2021 for the observational study.

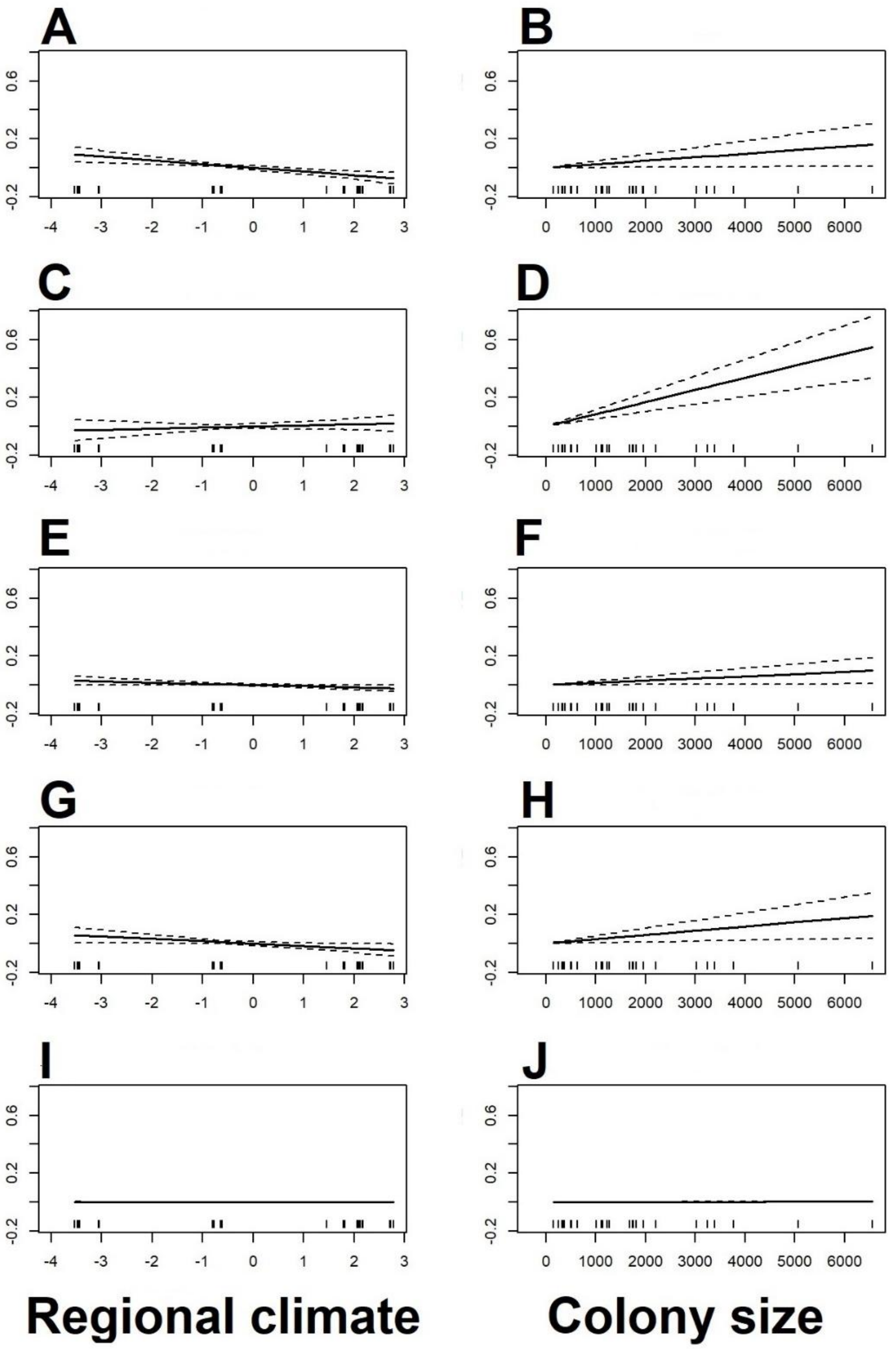


**Supplementary Figure 6.** Partial residuals plots of a multivariate normal additive model of the colony traits for the observational study according to colony size (total number of workers) and regional climate (PC1, positive values represent a colder and drier climate, while negative values represent a warmer and more humid climate) ($n$ = 26). (A, B)

Proportion of major workers; (C, D) Major worker mean body size (mm); (E, F) Worker body size variance ($mm^2$); (G, H) Worker head width variance ($mm^2$); (I, J) Worker head-to-body ratio variance ($mm^2$).

**Supplementary Table 1.** Principal component analysis of the bioclimatic variables expected to affect worker polymorphism.

| Bioclimatic variables | PC1 loading | PC2 loading |
|---|---|---|
| Mean diurnal range | -0.178 | 0.684 |
| Isothermality | -0.344 | 0.390 |
| Maximum temperature of warmest month | -0.377 | 0.224 |
| Mean annual temperature | -0.400 | 0.003 |
| Minimum temperature of coldest month | -0.391 | -0.174 |
| Annual precipitation | -0.390 | -0.118 |
| Seasonality | 0.386 | 0.197 |
| Temperature annual range | 0.305 | 0.496 |

**Supplementary Table 2.** Generalized additive models of five colony-level traits describing worker polymorphism according to biotic and abiotic predictors ($n$ = 26). Beta family was used for the proportion of major workers ($z$-values) and gaussian for the rest of the traits ($t$-values). We only kept the significant predictors ($p < 0.05$) for further analyses. All significant terms are shown in bold.

| Trait | Predictor | Estimate | Standard error | $z/t$-value | $p$ |
|---|---|---|---|---|---|
| Major worker proportion | Intercept | $-1.7\times10^{-1}$ | $5.4\times10^{-1}$ | -3.1 | 0.002 |
| | Regional climate | $-0.2\times10^{-1}$ | $5.8\times10^{-2}$ | -2.8 | 0.005 |
| | Local climate | $-1.0\times10^{-3}$ | $4.9\times10^{-3}$ | -0.2 | 0.821 |
| | Nest diameter | $-7.2\times10^{-3}$ | $5.7\times10^{-2}$ | -0.1 | 0.900 |
| | Colony size | $1.4\times10^{-4}$ | $1.0\times10^{-4}$ | 1.4 | 0.173 |
| | Competition | $-0.2\times10^{-1}$ | $4.0\times10^{-1}$ | -0.6 | 0.571 |
| Mean major worker size | Intercept | 3.3 | $1.2\times10^{-1}$ | 27.9 | < 0.001 |
| | Regional climate | $7.6\times10^{-3}$ | $1.2\times10^{-2}$ | 0.6 | 0.542 |
| | Local climate | $1.3\times10^{-3}$ | $1.1\times10^{-3}$ | 1.2 | 0.233 |
| | Nest diameter | $4.3\times10^{-4}$ | $1.3\times10^{-2}$ | 0.03 | 0.974 |
| | Colony size | $8.0\times10^{-5}$ | $2.2\times10^{-5}$ | 3.6 | 0.002 |
| | Competition | $-1.6\times10^{-1}$ | $7.7\times10^{-2}$ | -2.0 | 0.057 |
| Wor ker | Intercept | $1.2\times10^{-1}$ | $5.6\times10^{-2}$ | 2.1 | 0.045 |

| | | | | | |
|---|---|---|---|---|---|
| | Regional climate | $-1.1\times10^{-2}$ | $5.8\times10^{-3}$ | -2.0 | 0.066 |
| | Local climate | $4.7\times10^{-4}$ | $5.0\times10^{-4}$ | 1.0 | 0.353 |
| | Nest diameter | $-2.3\times10^{-3}$ | $6.0\times10^{-3}$ | -0.4 | 0.705 |
| | Colony size | $2.0\times10^{-5}$ | $1.0\times10^{-5}$ | 2.0 | 0.064 |
| | Competition | $-1.7\times10^{-2}$ | $3.6\times10^{-2}$ | -0.5 | 0.641 |
| Worker head size variance | Intercept | $1.8\times10^{-1}$ | $9.4\times10^{-2}$ | 1.9 | 0.071 |
| | Regional climate | $-2.0\times10^{-2}$ | $9.8\times10^{-3}$ | -2.1 | 0.053 |
| | Local climate | $7.1\times10^{-4}$ | $8.4\times10^{-4}$ | 0.8 | 0.409 |
| | Nest diameter | $-4.1\times10^{-3}$ | $1.0\times10^{-2}$ | -0.4 | 0.687 |
| | Colony size | $3.5\times10^{-5}$ | $1.8\times10^{-5}$ | 2.0 | 0.059 |
| | Competition | $-5.1\times10^{-2}$ | $6.2\times10^{-2}$ | -0.8 | 0.417 |
| Head-to-body ratio variance | Intercept | $4.7\times10^{-3}$ | $2.0\times10^{-3}$ | 2.3 | 0.031 |
| | Regional climate | $-4.4\times10^{-4}$ | $2.1\times10^{-4}$ | -2.1 | 0.047 |
| | Local climate | $1.6\times10^{-5}$ | $1.8\times10^{-5}$ | 0.9 | 0.393 |
| | Nest diameter | $-5.4\times10^{-5}$ | $2.1\times10^{-4}$ | -0.3 | 0.805 |
| | Colony size | $5.5\times10^{-7}$ | $3.7\times10^{-7}$ | 1.5 | 0.154 |
| | Competition | $-6.7\times10^{-4}$ | $1.3\times10^{-3}$ | -0.5 | 0.615 |

**Supplementary Table 3.** Comparison of three metrics of morphometric variation using generalized additive models of the standard deviation, coefficient of variance and variance of worker body size, worker head size and head-to-body ratio according to biotic and abiotic predictors using gaussian family (*n* = 26). All significant terms are shown in bold.

| | | Standard deviation | | Coefficient of variation | | Variance | |
|---|---|---|---|---|---|---|---|
| Trait | Predictor | Estimate | *p* | Estimate | *p* | Estimate | *p* |
| Worker body size | Intercept | $3.4\times10^{-1}$ | **< 0.001** | $1.3\times10^{1}$ | **< 0.001** | $1.2\times10^{-1}$ | **0.045** |
| | Regional climate | $-1.2\times10^{-2}$ | 0.091 | $-3.0\times10^{-1}$ | 0.178 | $-1.1\times10^{-2}$ | 0.066 |
| | Local climate | $5.0\times10^{-4}$ | 0.414 | $1.0\times10^{-2}$ | 0.517 | $4.7\times10^{-4}$ | 0.353 |
| | Nest diameter | $-2.5\times10^{-3}$ | 0.729 | $-1.0\times10^{-1}$ | 0.614 | $-2.3\times10^{-3}$ | 0.705 |
| | Colony size | $2.6\times10^{-5}$ | 0.055 | $1.0\times10^{-3}$ | 0.170 | $2.0\times10^{-5}$ | 0.064 |
| | Competition | $-1.5\times10^{-2}$ | 0.741 | $9.3\times10^{-1}$ | 0.525 | $-1.7\times10^{-2}$ | 0.641 |
| Worker head size | Intercept | $4.4\times10^{-1}$ | **< 0.001** | $2.4\times10^{1}$ | **< 0.001** | $1.8\times10^{-1}$ | 0.071 |
| | Regional climate | $-1.8\times10^{-2}$ | 0.072 | $-4.5\times10^{-1}$ | 0.279 | $-2.0\times10^{-2}$ | 0.053 |
| | Local climate | $3.8\times10^{-4}$ | 0.493 | $3.1\times10^{-2}$ | 0.381 | $7.1\times10^{-4}$ | 0.409 |
| | Nest diameter | $-5.5\times10^{-3}$ | 0.707 | $-1.2\times10^{-1}$ | 0.783 | $-4.1\times10^{-3}$ | 0.687 |
| | Colony size | $4.0\times10^{-5}$ | 0.050 | $9.9\times10^{-4}$ | 0.182 | $3.5\times10^{-5}$ | 0.059 |

| | | | | | | | |
|---|---|---|---|---|---|---|---|
| | Competition | $-2.5\times10^{-2}$ | 0.477 | $1.3\times10^{-1}$ | 0.958 | $-5.1\times10^{-2}$ | 0.417 |
| Head-to-body ratio | Intercept | $6.8\times10^{-2}$ | **< 0.001** | 9.8 | **< 0.001** | $4.7\times10^{-3}$ | **0.031** |
| | Regional climate | $-2.5\times10^{-3}$ | 0.060 | $-2.9\times10^{-1}$ | 0.089 | $-4.4\times10^{-4}$ | **0.047** |
| | Local climate | $8.2\times10^{-5}$ | 0.463 | $1.4\times10^{-2}$ | 0.335 | $1.6\times10^{-5}$ | 0.393 |
| | Nest diameter | $-3.3\times10^{-4}$ | 0.807 | $-1.3\times10^{-2}$ | 0.937 | $-5.4\times10^{-5}$ | 0.805 |
| | Colony size | $3.5\times10^{-6}$ | 0.136 | $4.7\times10^{-4}$ | 0.120 | $5.5\times10^{-7}$ | 0.154 |
| | Competition | $-2.9\times10^{-3}$ | 0.717 | $-2.2\times10^{-1}$ | 0.835 | $-6.7\times10^{-4}$ | 0.615 |

**Supplementary Table 4.** Multivariate normal additive model of five colony-level traits describing worker polymorphism according to regional climate and colony size as predictors. All significant terms are shown in bold.

| Trait | Predictor | Estimate | Standard error | *z*-value | *p* |
|---|---|---|---|---|---|
| Major worker proportion | Intercept | $1.3\times10^{-1}$ | $2.7\times10^{-2}$ | 4.8 | **< 0.001** |
| | Regional climate | $-2.6\times10^{-2}$ | $7.1\times10^{-3}$ | -3.7 | **< 0.001** |
| | Colony size | $2.4\times10^{-5}$ | $1.1\times10^{-5}$ | 2.2 | **0.031** |
| Mean major worker body size | Intercept | 3.4 | $3.9\times10^{-2}$ | 87.5 | **< 0.001** |
| | Regional climate | $7.4\times10^{-3}$ | $1.0\times10^{-2}$ | 0.7 | 0.473 |
| | Colony size | $8.4\times10^{-5}$ | $1.6\times10^{-5}$ | 5.1 | **< 0.001** |
| Worker body size variance | Intercept | $1.4\times10^{-1}$ | $1.7\times10^{-2}$ | 8.4 | **< 0.001** |
| | Regional climate | $-8.3\times10^{-3}$ | $4.4\times10^{-3}$ | -1.9 | 0.061 |
| | Colony size | $1.6\times10^{-5}$ | $7.0\times10^{-6}$ | 2.3 | **0.024** |
| Worker head size variance | Intercept | $2.0\times10^{-1}$ | $2.8\times10^{-2}$ | 7.0 | **< 0.001** |
| | Regional climate | $-1.6\times10^{-2}$ | $7.5\times10^{-3}$ | -2.2 | **0.029** |
| | Colony size | $2.9\times10^{-5}$ | $1.2\times10^{-5}$ | 2.5 | **0.013** |
| Head-to-body ratio variance | Intercept | $5.3\times10^{-3}$ | $5.9\times10^{-4}$ | 9.1 | **< 0.001** |
| | Regional climate | $-3.5\times10^{-4}$ | $1.6\times10^{-4}$ | -2.2 | **0.026** |
| | Colony size | $4.3\times10^{-7}$ | $2.5\times10^{-7}$ | 1.7 | 0.082 |

| |
|---|
| Deviance explained = 30.4%, -REML = -372.62, Scale estimate = 1, $n$ = 26 |

**Supplementary Table 5.** Generalized additive models of the proportion of major workers according to abiotic predictors, including number of days when temperature was above 0°C (colony activity) and 9°C (larval development) as well as regional mean annual temperature and regional mean diurnal range for the years 1970 to 2000 [31] ($n$ = 18). Colony size was included in each model. Beta family was used for the proportion of major workers. All significant terms are shown in bold.

| Predictor | Estimate | Standard error | $z$-value | $p$ | AICc |
|---|---|---|---|---|---|
| **Intercept** | -3.6 | $6.0 \times 10^{-1}$ | -6.0 | **< 0.001** | -34.35 |
| **Colony activity** | $9.5 \times 10^{-3}$ | $2.7 \times 10^{-3}$ | 3.5 | **< 0.001** | |
| **Colony size** | $1.5 \times 10^{-4}$ | $7.3 \times 10^{-5}$ | 2.1 | **0.038** | |
| **Intercept** | -3.2 | $6.0 \times 10^{-1}$ | -5.4 | **< 0.001** | -32.72 |
| **Larval development** | $1.6 \times 10^{-2}$ | $5.8 \times 10^{-3}$ | 2.8 | **0.005** | |
| **Colony size** | $1.8 \times 10^{-4}$ | $7.8 \times 10^{-5}$ | 2.3 | **0.024** | |
| **Intercept** | -1.9 | $2.4 \times 10^{-1}$ | -8.0 | **< 0.001** | -30.20 |
| **Regional mean annual temperature** | $1.3 \times 10^{-1}$ | $5.5 \times 10^{-2}$ | 2.3 | **0.022** | |
| Colony size | $1.5 \times 10^{-4}$ | $8.1 \times 10^{-5}$ | 1.9 | 0.063 | |
| **Intercept** | -3.6 | 1.7 | -2.1 | **0.036** | -26.30 |
| Regional mean diurnal range | $1.9 \times 10^{-1}$ | $1.7 \times 10^{-1}$ | 1.1 | 0.252 | |

| | | | | |
|---|---|---|---|---|
| Colony size | $5.7\times10^{-5}$ | $9.3\times10^{-5}$ | 0.6 | 0.539 |

**Supplementary Table 6.** Number of *C. herculeanus* colonies where competitive species workers occurred within 10 meters around their nest ($n$ = 26). We considered *Formica* strong competitors.

| Species | Occurrence |
|---|---|
| *Camponotus nearcticus* Emery, 1893 | 1 |
| *Dolichoderus mariae* Forel, 1885 | 1 |
| *Dolichoderus pustulatus* Mayr, 1886 | 1 |
| *Formica neorufibarbis* Emery, 1893 | 4 |
| *Formica subaenescens* Emery, 1893 | 1 |
| *Lasius americanus* Emery, 1893 | 1 |
| *Leptothorax* spp. Mayr, 1855 | 11 |
| *Myrmica alaskensis* Wheeler, 1917 | 10 |
| *Myrmica fracticornis* Forel, 1901 | 2 |
| *Myrmica lobifrons* Pergande, 1900 | 1 |
| *Tapinoma sessile* Say, 1836 | 2 |
